# The *C. elegans* WASH complex supports epithelial polarity by promoting endosomal sorting of Cadherin

**DOI:** 10.64898/2025.12.23.696084

**Authors:** Patricia Irizarry-Barreto, Jennifer Smolyn, Victoria Brown, Bhagavathi Ramamurthy, Martha C. Soto

## Abstract

Epithelial polarity requires polarized distribution of the apical adhesion complex that connects cells through the Cadherin transmembrane protein. Cadherin is intimately linked to the actin cytoskeleton through alpha catenin, which directly binds F-actin to set up the apical actin belt. Branched actin is formed when the Arp2/3 complex is activated by Nucleation Promoting Factors. *C. elegans* has three Nucleation Promoting Factors, WASP, WAVE and WASH. Our studies showed that WAVE-dependent branched actin promotes apical transport of Cadherin, including apically-directed transport of RAB-11-enriched endosomes. However, the contribution of other Nucleation Promoting Factors to Cadherin polarity has not been examined. The *C. elegans* WASH complex is not well described. Here we characterize components of the WASH complex, and confirm that CO5G5.2, despite being highly divergent, is the functional FAM21 component in *C. elegans*. We show that the WASH complex is enriched at early and recycling endosomes in the adult intestine, where it supports retrograde Cadherin transport. Our findings demonstrate that individual branched actin regulators promote specific transport steps, and identify WASH function at RME-1/EHD-enriched endosomes as an important contributor to Cadherin polarity and cargo sorting in a mature epithelium.

**Significance Statement:** \**C. elegans* has three known NPF (nucleation promoting factor) complexes to activate Arp2/3: WAVE, WASP and WASH. However, the role of *C. elegans* WASH complex in intracellular transport was unreported, and the essential component, FAM21, was not identified.

*We show that *C. elegans* C05G5.2 is a likely FAM21 ortholog, since it functions in collaboration with other WASH components in retrograde transport. Loss of WASH components reduced epithelial polarity, while the cargo Cadherin was mis-sorted from early endosomes.

*WASH branched actin polarizes epithelia using a functional FAM21 ortholog to promote sorting of retrograde cargo, including Cadherin.

## INTRODUCTION

*C. elegans* has three Nucleation Promoting Factors (NPFs) WASP, WAVE, and WASH, that can activate the Arp2/3 complex, promoting branched actin formation. Branched actin promotes movements and dynamics of cellular events, therefore it plays a large role in protein transport. The NPF WASP supports endocytosis in yeast, mammals, and *C. elegans* (Engqvist-Goldstein & Drubin 2003; Benesch et al., 2005; Giuliani et al., 2009). Our work showed a role for WAVE in transport (Guiliani et al., 2009; Patel et al., 2013), including apically directed RAB-11 transport, and WAVE accumulation at the Golgi (Cordova-Burgos et al., 2023). The discovery of WASH was accompanied by findings that it has an important role at endosomes (Derivery et al., 2009; Gomez and Billadeau 2009; Romano-Moreno et al., 2024). WASH is unique among the highly conserved Arp2/3 NPFs, since it directly links membrane dynamics with actin and microtubule (MT) cytoskeletons (Fig. 1A,B). In *Drosophila*, WASH bundled MTs, but WASP and SCAR/WAVE could not (Liu et al., 2009). Since invertebrates like *C. elegans, Dictyostelium* and *Drosophila* do not have orthologs of the NPF WHAMM (Campellone et al., 2008), WASH may work in invertebrates to sort proteins at Golgi-ER (Gomez & Billadeau 2009).

**Figure 1.**
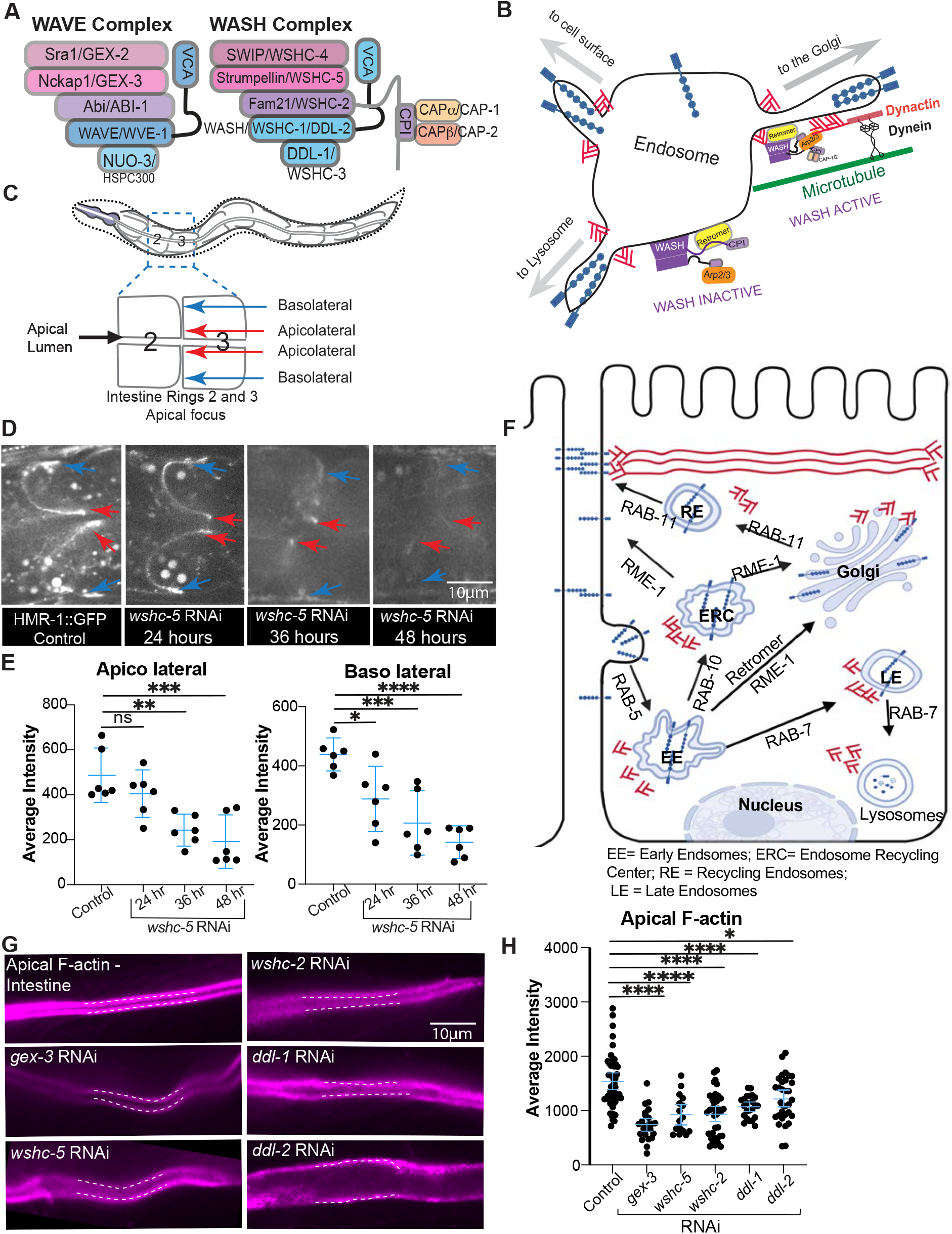
*C. elegans* WASH complex supports Cadherin enrichment and apical polarity. (A) The WAVE and WASH complexes, which promote branched actin as Nucleation Promoting Factors (NPFs) for Arp2/3, have similar structure. WAVE and WASH are composed of five proteins, including the paralogs WAVE/WVE-1 and WASH/WSHC-1/DDL-2, which activate Arp2/3 with their VCA (verprolin central acidic) domains. (B) Model of WASH complex (purple) on an endosome highlighting its interactions with Retromer (yellow) at the endosomal membranes and with the Dynactin/Dynein complex, on microtubules. The interaction with Dynactin activates WASH-dependent branched actin that is proposed to support vesicle tubulation and scission (see also 1F). (C) Cartoon of *C. elegans* intestine with a focus on Rings 2 and 3, composed of two cells each, where most of the measurements were done. Rings 2/3 are behind the pharynx and before the germline, making live imaging simple. Most images are shown from an apical focus, which allows comparison of protein enrichment at apical lumen, apicolateral (red arrows), and basolateral (blue arrows) membrane regions. (D,E) RNAi depletion of *wshc-5* was monitored for effects on Cadherin/HMR-1::GFP. (F) Model for how WASH-dependent branched actin may support HMR-1/Cadherin (blue) transport in epithelia. WASH is predicted to support retromer transport, from early endosomes (EE) to the Golgi. (G,H) Intestinal apical F-actin was measured using strain OX966 *Pglo-1-::Lifeact-Tag-RFP*, in Controls and animals depleted of *gex-3, wshc-5, wshc-2, ddl-1* and *ddl-2* using RNAi for 48 hours. Statistical analyses here and elsewhere used one-way ANOVA, unless stated otherwise. Asterisks here and in all figures mark statistical significance: \**p* < 0.05, \*\**p* < 0.001, \*\*\**p* < 0.0001, \*\*\*\**p* < 0.00001. Error bars show 95% confidence intervals in all figures.

The least studied of the three NPFs in *C. elegans* is the WASH complex. Since the five components of the WASH complex have multiple names across species, we refer to the *C. elegans* WASH complex orthologs by their WSHC1-5 name and where it differs, also by their *C. elegans* name. Two components of the *C. elegans* WASH complex, WSHC-1/*ddl-2*/WASH and *WSHC-3/ddl-1*/CCDC53, were named “*ddl*” as *<u>d</u>af-16*-<u>d</u>ependent longevity genes in a genome-wide RNAi screen (Hansen et al., 2005). WSHC-1/DDL-2 and WSHC-3/DDL-1 were shown to regulate heat-shock transcription factor HSF-1 to modulate longevity and thermotolerance (Chiang et a., 2012). A third WASH component, WSHC-5/Strumpellin, was CRISPR tagged at its endogenous locus, and shown to localize in puncta in embryos (Zhang et al., 2025). Our research group conducted the first characterization of WASH in *C. elegans* transport, and suggested it supports retrograde transport (Smolyn Master’s Thesis, 2020).

Protein transport of adhesion molecules is essential for the formation and maintenance of healthy junctions (Bruser and Bogdan, 2017). Cadherin transport was proposed to require actin (Woichansky et al., 2016), but the actin regulators are only recently identified, including WAVE that supports apically directed transport on RAB-11 endosomes (Cordova-Burgos et al., 2023). WAVE also supports endosomal transport of yolk proteins (Guiliani et al., 2009) and transport of Wls/MIG-14, TGN-38 and Cadherin retrograde recycling from recycling endosomes to the Golgi (Bai and Grant 2015; Cordova-Burgos et al., 2023). The role of WASH in Cadherin transport has not been investigated.

## RESULTS and DISCUSSION

### *C. elegans* WASH complex supports apical enrichment of Cadherin

Our previous studies demonstrated that WAVE supported Cadherin transport. However, WAVE was not enriched at all endosomal organelles and did not alter Cadherin enrichment at all endosomal organelles, suggesting other branched actin regulators may be involved. We previously found that loss of WASP had minor effects on epithelial transport (Patel et al., 2013). However, WASH, a protein complex with striking similarities to the WAVE complex (Fig. 1A,B), had not been examined for effects on Cadherin.

We monitored the distribution of Cadherin, a transmembrane protein that is transported with the help of WAVE branched actin, but which has not been examined for transport by WASH. Endogenously tagged E-Cadherin/HMR-1::GFP (Marston DJ, et al. 2016) is normally enriched at the apicolateral regions of epithelia, like adult intestine, with lower enrichment at more basolateral regions (Cordova-Burgos 2020, 2023, Fig. 1C). To test if Cadherin enrichment depended on WASH, we depleted WASH component *wshc-5* using RNAi (as in Cordova-Burgos et al., 2023). Loss of any core WASH component (*wshc-2, wshc-5* or *wshc-4/SWIP*) is expected to destabilize the WASH complex (Jia et al., 2010). Depleting *wshc-5*/Strumpellin for 24 hours showed mild changes in Cadherin/HRMI-1::GFP levels, 36 hour depletion showed stronger changes, while 48 hour depletion strongly reduced Cadherin/HMR-1::GFP levels, making it difficult to image (Fig. 1D,E).

To directly visualize the consequences of Cadherin apicolateral loss, we imaged F-actin using Lifeact expressed in the intestine (Cordova-Burgos et al. 2021) and compared controls to animals depleted of *wshc-5* and WAVE component *gex-3* for 36 hours. As previously shown, loss of *gex-3* significantly decreased apical F-actin in the adult intestine. Loss of *wshc-5, wshc-2, ddl-1* or *ddl-2* also decreased apical F-actin. Thus, the decrease in Cadherin caused by depleted WASH complex has consequences for epithelial polarity of the adult intestine (Fig. 1G,H).

### WASH supports cellular organization of endosomal organelles

WASH is expected to be enriched at endosomes (Derivery et al., 2009; Gomez and Billadeau 2009; Romano-Moreno et al., 2024, Fig. 1F). To understand what occurs at distinct endosomal organelles when WASH components are depleted, we compared control animals with animals depleted of *wshc-5*, for the patterns of early endosomes enriched in RFP::RAB-5, late endosomes enriched for RFP::RAB-7, recycling endosomes enriched for RFP::RME-1 or RFP::RAB-10, and the Golgi, where AMAN-2::GFP is enriched (Treusch *et al*., 2004; Gleason *et al*., 2016).

Loss of *wshc-5* increased the intensity, and vesicle size of endosomes enriched for RFP::RAB-5, which includes early endosomes (Fig. 2A,B-D see also Fig. 1F). Loss of *wshc-5* also increased the intensity and number of RFP::RAB-7 enriched endosomes, which include maturing late endosomes (Fig. 2A,B-E). By contrast, loss of *wshc-5* resulted in RFP::RME-1-positive recycling endosomes with decreased intensity and decreased vesicle numbers. RFP::RME-1 is most enriched in basolateral regions and those regions showed the largest changes. RFP::RAB-10 is normally found throughout the cytoplasm. Depletion of *wshc-5* reduced the intensity and size of endosomes enriched for RFP::RAB-10, in all regions of the intestinal cells (Fig. 2A,B-D). GFP::RAB-11 is normally highly enriched at apical regions. Depleting *wshc-5* resulted in reduced overall signal, most visible at apical regions. Finally, the Golgi protein AMAN-2::GFP is found in small bright puncta, the Golgi ministacks, all throughout the intestine. Depletion of *wshc-5* resulted in increased intensity of AMAN-2::GFP, most visible at basal regions. Collectively, these data suggest an overall shift in endosomal organization, with increased intensity of proteins enriched at early endosomes (RFP::RAB-5), late endosomes (RFP::RAB-7) and the Golgi (GFP::AMAN-2), at the expense of proteins enriched at recycling endosomes (RFP::RME-1, RFP::RAB-10 and GFP::RAB-11) (Fig. 2C-E).

**Figure 2.**
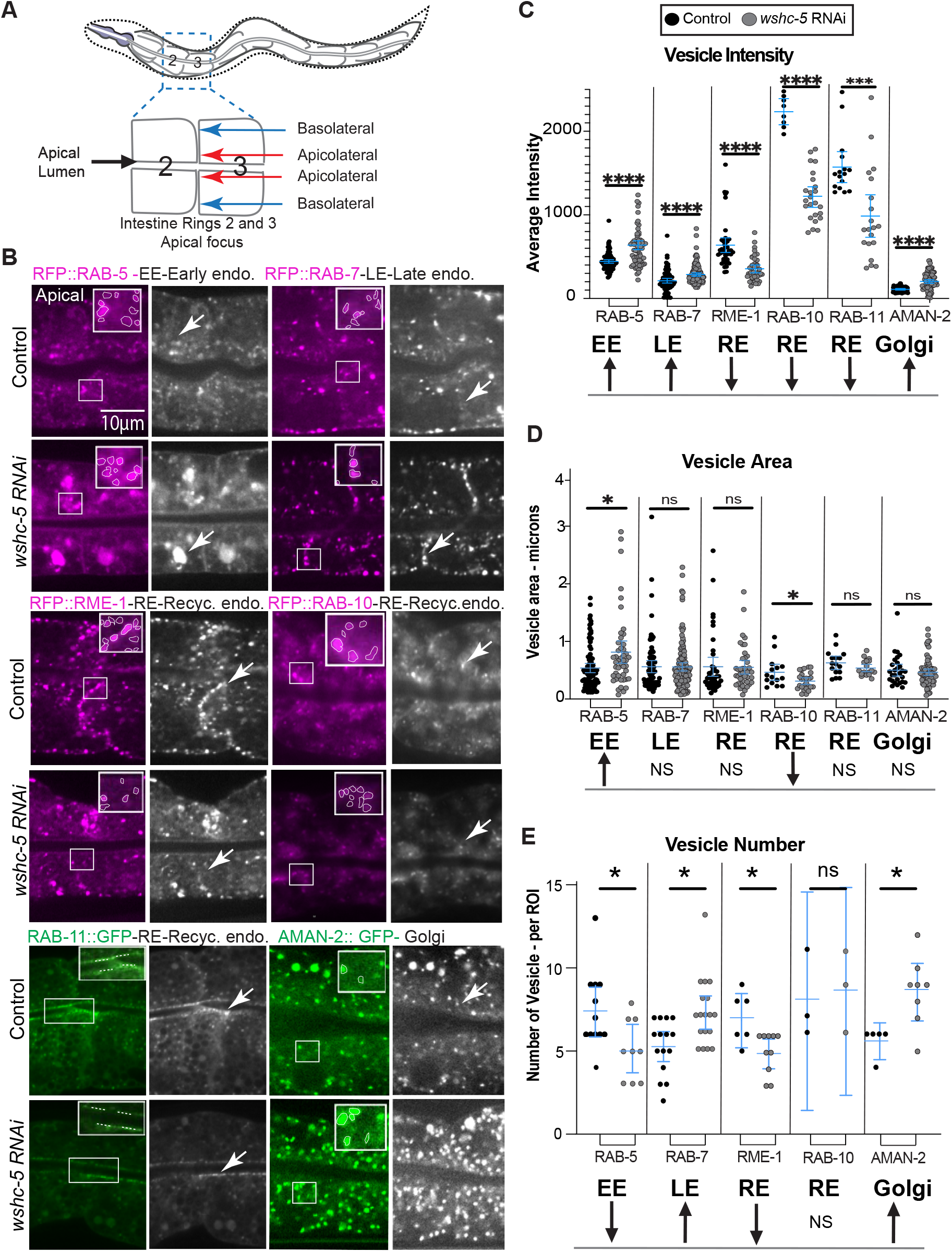
WASH supports cellular organization of endosomal organelles. (A). Cartoon of *C. elegans* intestinal region used for imaging and analysis. (B) Apical views of intestinal Rings 2/3 are shown for each endosomal organelle. Top images are Controls and immediately below are *wshc-5*-depleted animals. The white box indicates the region that is enlarged to the right, and the region were measurements were made. (C,D,E) The white freehand curves or lines at puncta shown in (B) were compared for intensity, vesicle area, and vesicle number, in equally sized regions of individual worms. 3-8 animals were compared for each genotype.

Some cargoes travel from early endosomes, with the help of RAB-10 enriched endosomes, to endosomes enriched for RME-1 for further sorting (Chen et al., 2006). These findings could indicate a requirement for WASH at early endosomes for the sorting of cargos normally destined for recycling endosomes. These findings thus encouraged us to investigate where WASH is enriched in the endosomes of the intestine.

### WASH is enriched at some endosomal organelles

To determine WASH subcellular enrichment in intestinal epithelia, we used endogenously tagged mNG::WSHC-5, previously localized to puncta in embryos (Zhang et al., 2025; Fig. 3A). In adult intestines, we noted mNG::WSHC-5 also enriched at puncta (Fig. 3A). Since intestines have autofluorescence produced by the lysosomal gut granules (Morris et al., 2018), we subtracted the autofluorescence signal before analysis (Fig. 3A, Fig. S1, and Methods).

**Figure 3.**
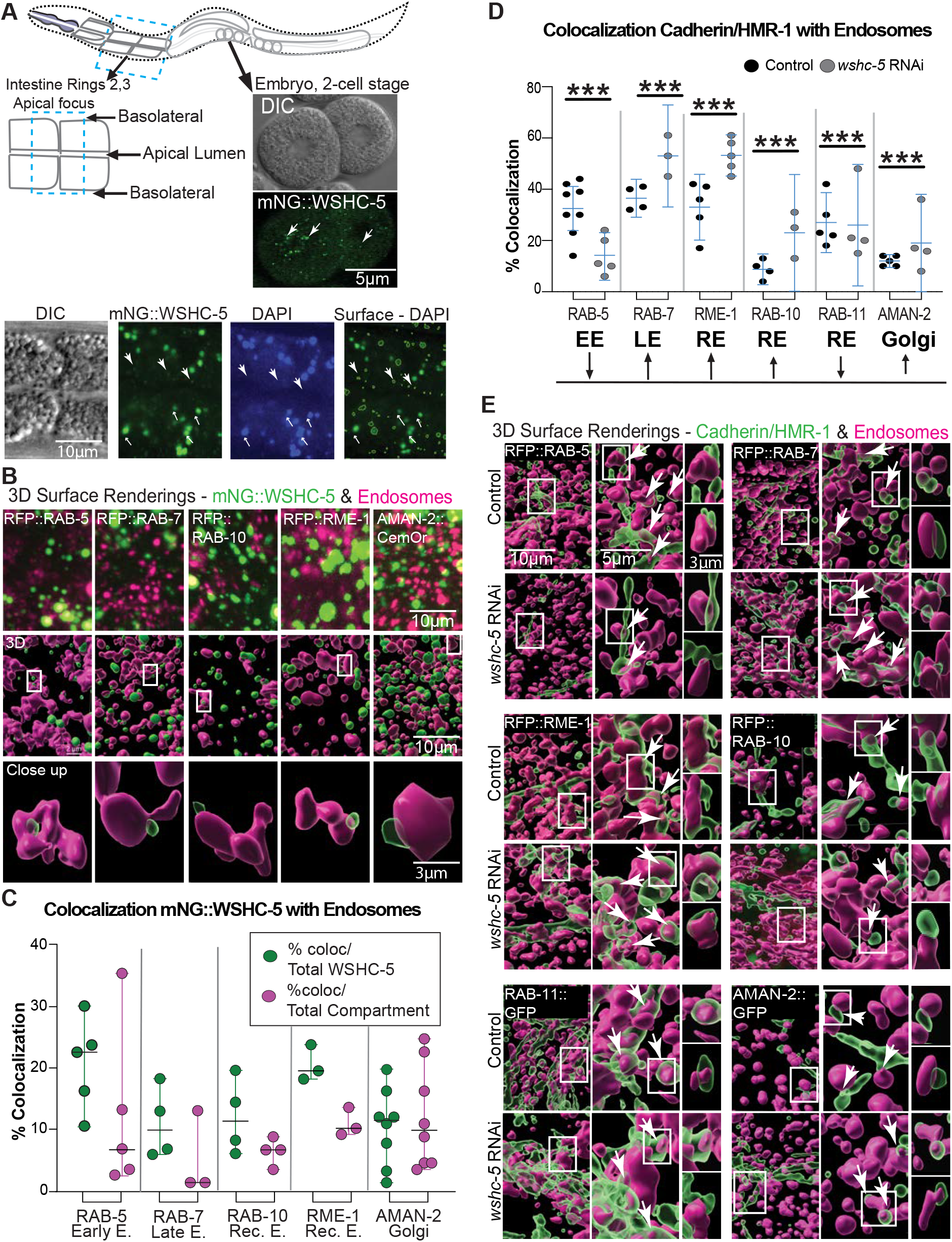
WASH is enriched at recycling endosomes and supports Cadherin transport through recycling endosomes. (A) Cartoon worm highlights intestinal region, Rings 2/3, and germline. The expression of mNG::WSCH-5 is shown in a two-cell embryo (top), and in an L4 intestine (below). Intestinal autofluorescence (DAPI, blue channel) was subtracted (right panel) to better show mNG::WSHC-5 intestinal enrichment. Large arrows indicate mNG::WSHC-5 puncta. Small arrows indicate large autofluorescent gut granules. (B) *Top row:* Representative raw data 3D projections of Ring 2/3 region of intestine, prepared in Fiji, showing Maximum Projections, with mNG::WSCH-5 in Cyan and endosomes in Magenta. *Middle Row:* 3D Surface renderings of intestine Ring 2/3 region, full width of a worm, done in IMARIS, to measure colocalization. *Bottom row:* Close Up to illustrate colocalization. The green surfaces are shown partially transparent to better show the overlap. See also Fig. 1S to view how the raw data was used to create the 3D surface renderings in IMARIS. (C) Colocalization of mNG::WSHC-5 (green) and endosomal organelles (magenta). Green dots report % endosomes colocalizing with total mNG::WSHC-5. Magenta dots report % mNG::WSHC-5 colocalizing with total endosomes. Each dot is the average from a separate experiment, based on hundreds of measurements done in IMARIS (see Methods, Fig. S1). N = at least 3 experiments per measurement. (D) Colocalization of HMR-1/Cadherin with the endosomal organelles was compared in Controls and animals depleted of WSHC-5 via RNAi for 36 hours. Colocalization was measured using IMARIS software as in 3B,C. After creating the surfaces using the object-to-object tool, we measured the distance between the surfaces created in the green and the red channel. Zero distance between surfaces was counted as colocalization. DAPI was subtracted to remove autofluorescent signal from lysosomal organelles. (E) Controls and animals depleted of WSHC-5 for 36 hours, are shown from the Apical focus at Rings 2/3, to show overlap of the surfaces in the green (HMR-1) and magenta (endosomes) channels. Arrows point to colocalization regions. White box = area selected for close up to the right. Close ups at the far right are two views of one region of overlap, to better show colocalization from different angles.

To test if the WSHC-5 puncta are enriched at specific endosomes or the Golgi, we crossed mNG::WSHC-5 into marker strains tagged with RFP or CemOrange2 (Thomas et al., 2019; Cordova-Burgos et al., 2023). 3D projections of the raw data suggested partial overlap (Fig. 3B). Using simultaneous capture imaging, 3D rendering, and machine learning approaches to distinguish signals from autofluorescence, we measured colocalization based on overlap volume ratios and nearest neighbor analysis. WSHC-5 was most enriched at early endosomes (23%), followed by RME-1-positive recycling endosomes (20%), RAB-10-positive recycling endosomes (12%), RAB-7-positive late endosomes (10%), and at similar levels at the Golgi (10%) (Fig. 3C). Thus, a significant amount of WASH is enriched at early endosomes and recycling endosomes, which supported a role in transport.

We next examined which endosomes are enriched for Cadherin accumulation at steady state in the adult intestinal endothelia by crossing endogenously tagged Cadherin, either HMR-1::mKate2 or HMR-1::GFP, into the endosomal strains used for Fig. 3B,C. Using a similar approach as in Fig. 3C, we quantified Cadherin/HMR-1::GFP enrichment at different endosomal organelles. Cadherin was found enriched at 28% of RFP::RAB-5 early endosomes, 36% of RFP::RAB-7 late endosomes and 32% of RFP::RME-1 recycling endosomes, 8% of RFP:: RAB-10 endosomes, 28% of RAB-11::GFP endosomes and 12% of AMAN-2::GFP Golgi (Fig. 3D,E).

Depleting WASH with *wshc-5* RNAi for 36 hours resulted in reduced levels of Cadherin overall, and a shift in the distribution of Cadherin. Cadherin enrichment at RFP:: RAB-5 early endosomes dropped from 28% to 14%, while RAB-7::RFP late endosomes increased from 36% to 53% (Fig. 3D). There was increased Cadherin enrichment at RFP::RME-1 recycling endosomes, from 32% to 51%. This suggests that Cadherin transport from early endosome and recycling endosomes requires the WASH complex. With reduced WASH complex, Cadherin sorting changed, and more accumulated at late endosomes and RME-1-positive recycling endosomes. Decreased Cadherin levels seen with depleted WASH are likely due to increased lysosomal degradation, since we showed that blocking lysosomal degradation in animals depleted of WASH, using the *cup-5* mutation, restored the levels of other cargos including the aquaporins AQP-1::GFP and AQP-4::GFP (Smolyn thesis, 2020). The increased Cadherin accumulation seen with depleted WASH at RME-1-positive endosomes was surprising since those endosomes also showed reduced intensity and number (Fig. 2).

### *C. elegans* possesses a complete pentameric WASH complex

To further address the role of *C. elegans* WASH in the transport of polarized cargos, we needed to identify the expected five components of *C. elegans* WASH complex (Fig. 1A). One problem was the apparent lack of a WSHC-2/FAM21 ortholog. WSHC-2/FAM21 function is central to WASH complex function in other species, since its long C-terminal tail connects WASH to two types of partner molecules at endosomes: Retromer components (VPS-29, VPS-35, VPS-26) and the Capping Proteins (CapZA/CAP-1 and CapZB/CAP-2) (Fig. 4A). Retromer promotes retrograde transport of proteins, while Capping Proteins are displaced when WASH binds to the Dynactin-Dynein complex during WASH activation (Figure 4A) (Seaman et al., 1998; Hierro et al., 2007; Fokin et al., 2021; Romano-Moreno et al., 2024). However, others have noted that FAM21 can be highly divergent. For example, Velle and colleagues showed that using a mutual-best-BLAST-hit approach, Drosophila did not appear to have a FAM21 ortholog (Velle et al., 2024).

**Figure 4.**
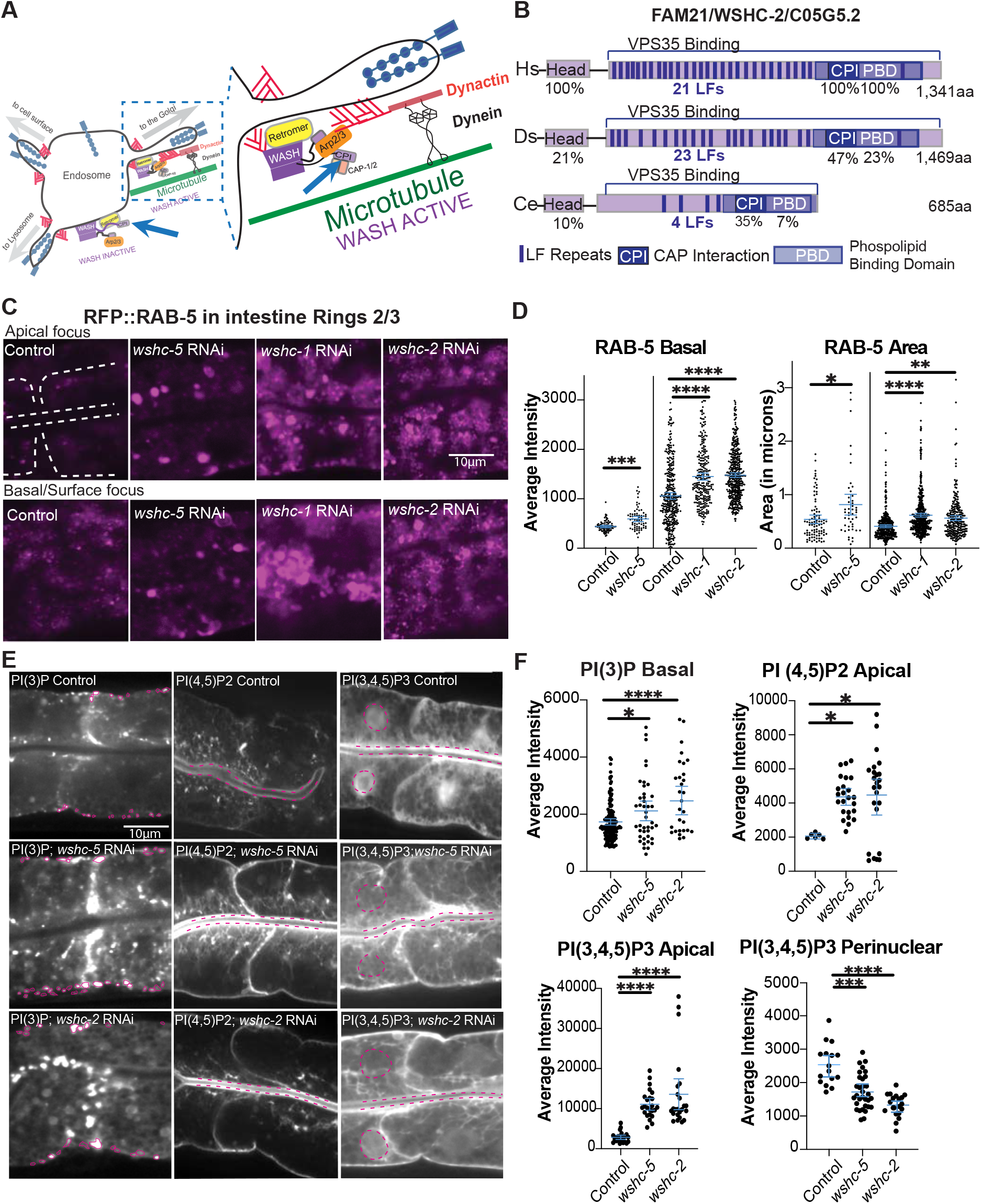
*C. elegans* C05G5.2 is a FAM-21/WSHC-2 ortholog with similar functions as other *C. elegans* WASH components. (A) Cartoon from Fig. 1B with enlarged area to highlight roles of WSHC-2/FAM21, the WASH component whose long C-terminal tail binds Retromer and removes CAPα/β/ CAP-1/2 from Dynactin during WASH activation. (B) The FAM21/WSHC-2 ortholog of *C. elegans*, C05H5.2, is highly divergent. Protein structure of *Caenorhabditis* elegans (Ce) C05G5.2 is aligned to FAM-21/WSHC-2 in *Homo* sapiens*(Hs)* and *Drosophila* melanogaster (Ds). Key domains, including LF repeats (dark purple), Cap Protein Interaction domain (CPI) and Phospholipid Binding Domain (PBD) are labeled with conserved identity listed. (C) Loss of WSHC-2/FAM21/CO5G5.2 is compared to loss of WASH components WSHC-5/Strumpellin and WSHC-1/DDL-2, using RNAi for 36 hours, at RFP::RAB-5 enriched early endosomes. (D) Quantification of vesicle intensity and vesicle area of RFP::RAB-5 endosomes, measured using the Freehand tool in Fiji to circle vesicles. (E) Effect of WSHC-5/Strumpellin and WSHC-2/FAM21/CO5G5.2 depletion, by RNAi for 48 hours, on phospholipid levels and distribution in the intestinal cells of *C. elegans*. The distribution of membrane lipids PI(3)P; PI(4,5)P2; and PI(3,4,5)P3 is shown from the Apical view. (F) Quantification of fluorescence intensity measured in FIJI using the line or freehand tool.

Aligning the highly divergent protein, CO5G5.2, with human and Drosophila WSHC-2/FAM21 revealed a likely reason this WASH component was initially missed (Figure 4B). The proposed *C. elegans* WSHC-2/FAM21 protein is about half the length of orthologs in other organisms. In addition, CO5G5.2 contains highly divergent WSHC-2/FAM21 motifs, as defined in Jia et al., 2010. Compared to human WSHC-2/FAM2, the N-terminal “head” domain (Jia et al., 2012), predicted to bind WSHC-1/DDL-2, showed only 10% identity over 220 residues, the Phospholipid binding domain that binds Retromer, showed 7% identity over 404 residues, and the Cap Protein Interacting (CPI) motif, that binds to CAPZ proteins showed 35% identity over 17 residues. Human WSHC-2/FAM21 has 21 LF (Leu-Phe) motifs, that connect WSHC-2 to Retromer components VPS35 and VPS29 (Jia et al. 2012; Romano-Moreno et al., 2024), *Drosophila* has 23, but *C. elegans* has only 4 LF motifs (Fig. 4B).

Structural work on human FAM21 suggested that while WSHC-2/FAM21 has 21 LF repeats, only a few bind to retromer. Only FAM21 Repeats 1-4 and 20, 21 are important for binding to retromer, and only 20 and 21 bind singly and strongly (Romano-Moreno et al, 2024). Thus, it is possible a WSHC-2/FAM21 ortholog with only 4 LF repeats could carry out WSHC-2/FAM21 retromer binding. In addition the 6/17 conserved residues in the CPI consensus motif (Jia et al. 2010, Hernandez-Valadares et al. 2010) may permit C05G5.5 to bind to the *C. elegans* Capping Proteins, CapZA/CAP-1 and CapZB/CAP-2 (Figure 4A,B), especially since three residues essential for CP binding (KxRxK) are conserved or similar (KxRxR) in WSHC-2/C05G5.2.

### The proposed FAM21 ortholog has a role in retromer transport

WSHC-2/FAM21 and the WASH complex regulate Retromer-dependent sorting (Gomez and Billadeau 2009). If the *C. elegans* WSHC-2 ortholog works with WASH to support retromer transport from early endosomes to the Golgi, loss of WSHC-2 was expected to alter the early endosomes (Fig. 3). In support of a Retromer function, loss of WSHC-2, similar to loss of WSHC-5 or WSHC-1/DDL-2 using RNAi depletion led to increased intensity and size of early endosomes enriched for RFP::RAB-5, (Fig. 4C), and as was shown for RAB-5 early endosome morphology in the absence of retromer components (Bai and Grant 2015 ).

### *C. elegans* WASH regulates membrane enrichment of phospholipids

Changes in endosome organization could result in changes in the phosphatidylinositol phospholipid (PIP) composition that establishes distinct cellular membranes. To further investigate the consequences of *wshc-2* depletion, we tested the effects of WSHC-2 and other WASH component depletion on phospholipids. RAB-5 recruits PI3 kinase to produce and enrich phosphatidylinositol 3-monophosphate (PI(3)P) on early endosomes. Depletion of WASH components *wshc-5* or *wshc-2*/CO5G5.2 for 48 hours significantly increased PI(3)P enrichment and intensity. This result supports the idea that CO5G5.2 behaves similarly as other WASH components in regulating PI(3)P enrichment. Two other phospholipids, Phosphatidylinositol-4,5-bisphosphate (PI(4,5)P2) and phosphatidylinositol-3,4,5-triphosphate (PI(3,4,5)P3), are normally enriched at the plasma membrane (Shi et al, 2012). RNAi depletion of *wshc-5* or CO05G5.2 increased plasma membrane enrichment of PI(4,5)P2 and PI(3,4,5)P3 (Fig. 4E). For PI(3,4,5)P3 we noted a significant drop in the perinuclear accumulation when all WASH components, including CO5G5.2 were depleted. While the molecular mechanisms that result in these phospholipid changes are not yet clear, these data demonstrated that CO5G5.2 loss phenocopies loss of other WASH components. Thus, the highly diverged WSHC-2/FAM21 ortholog CO5G5.2 is likely a component of the *C. elegans* WASH complex, and we refer to it hereafter as WSHC-2 (Figure 4F).

### Model for how WASH supports polarized transport

We propose that *C. elegans* has a highly diverged WSHC-2 ortholog that nevertheless contributes to WASH function. WASH in *C. elegans* appears to support retromer transport, as depleting WASH components leads to altered early endosomes, reduced WASH accumulation at recycling endosomes, and increased accumulation at late endosomes. One important epithelial cargo, Cadherin/HMR-1, depends on WASH for normal levels, in part due to a strong requirement for WASH at RME-1/EHD1 enriched endosomes. Loss of WASH components also affects sorting to recycling endosomes. Of note, loss of WASH had a distinct role on Cadherin transport, as compared to the effects of depleting WAVE, which did not affect early endosomes, but strongly affected RAB-11-positive recycling endosomes (Cordova-Burgos et al., 2023). These findings support the model that each Arp2/3 NPF has a distinct role in endocytosis and highlight the need to identify partner molecules that bring the distinct NPFs, WASH, WAVE and WASP, to distinct endosomal organelles to support sorting that then supports polarized distribution of cargos.

## Materials and methods

### C. elegans strains built for this paper

OX237 *mNG::wshc-5; gfp; tagRFP::rab-5;* OX1082 *mNG::wshc-5; tagRFP::rab-7;* OX1080 *mNG::wshc-5; tagRFP rab-10; OX1084 mNG::wshc-5; tagRFP::rme-1;* OX1083 m *mNG::wshc-5; aman-2::CemOr;* OX1081 *mNG::wshc-5; pGlo-1::LAmCh*.

### Strains used in this paper

LP902 *mNG::wshc-5* (Zhang et al., 2025); OX983 *hmr-1::mKate-2; aman2::gfp(Cordova-Burgos et al. 2023);* OX902 *hmr-1::gfp; tagRFP::rme-1(Cordova-Burgos et al. 2023);* OX810 *hmr-1::gfp; tagRFP::rab-5 (Cordova-Burgos et al. 2023);* OX806 *hmr-1::gfp; tagRFP::rab-7(Cordova-Burgos et al. 2023);* OX901 *hmr-1::gfp* ; *tagRFP rab-10 (Cordova-Burgos et al. 2023)*; OX971 *hmr-1::mKate2*; *gfp::rab-11 (Cordova-Burgos et al. 2023);* LP172 *hmr-1::gfp* (Marston et al., 2016); hmr-1::gfp; aman-2::cem-orange. *tagRFP::rab-5* (Gleason et al., 2016); *tagRFP::rab-10* (Gleason et al., 2016); *tagRFP::rab-7* (Gleason et al., 2016); *tagRFP::rme-1* (Gleason et al., 2016); RT311 *vha-6p::gfp::rab-11 (Chen et al, 20006);* OX966 *pGlo-1::TAG::RFP*; pRF4 rol-6(su1006), (Cordova-Burgos et al., 2021).

### RNAi experiments

All RNAi bacterial strains used in this study were administered by the feeding protocol as in (Sasidharan et al., 2018). RNAi feeding experiments were done at 23°C unless otherwise mentioned. Worms were synchronized and transferred onto seeded plate containing RNAi-expressing bacteria. To monitor effectiveness of the RNAi we used two methods: (1) We counted the percent dead embryos, which after two days is expected at >90% for *gex-3* and at least 10% for *wshc-5;* (2) We monitored post-embryonic silencing of a mNG-tagged strain in the intestine, LP902 *mNG::wshc-5*. Each Figure reports if RNAi treatments were done for 24, 36 or 48 hours. For example, Phospholipid strains were treated for 48 hours while organelle and Cadherin/HMR-1 strains were treated for 36 hours, so the Controls were monitored at the same time points.

### Live Imaging

Imaging was done in a temperature-controlled room set to 23°C on a Laser Spinning Disk Confocal Microscope with a Yokogawa scan head, on a Zeiss AxioImager Z1 Microscope using the Plan-Apo 63X/1.4NA oil lens. Imaging for colocalization studies used Visiview software and two identical Hamantsu CMOS Cameras to simultaneously capture two fluorophores. Each Figure explains imaging conditions, and only experiments done under the same conditions were compared. Images were analyzed using ImageJ or IMARIS, as explained in the Figure Legends. Controls and mutants were imaged within 3 days of each other with the same imaging conditions. All measurements were performed on raw data using ImageJ and/or IMARIS. Background intensity was subtracted by measuring average intensity in the same focal plane, near the animal.

### Microscopy of L4s and adults

Young adult stage animals (one day after L4 stage) and L4 larvae were placed on 10% agar pads in M9 solution and immobilized using 2μl of Levamizole(100μM) salts and covered with 1.5um coverslips. Images were taken within 15 minutes of making the pads. Imaging was done on a Zeiss AxioImager Z1 with a Yokogawa CSUX1-5000 spinning disc, using the Plan Apo 63X/1.4NA Oil lens.

### Quantitation of immunofluorescence

Quantitation of live fluorescence was performed using the line selection and the dynamic profile function of ImageJ to measure fluorescence along lines of equal length along, for example, the apical intestine or lateral regions of the intestinal cells. Some puncta were measured using the Freehand Tool to detect size changes. For all experiments shown, the images were captured at the same exposure settings for wild type and mutants. All quantitation was done on the raw images. The figure legends indicate when images were enhanced for contrast, and the same enhancement was applied to a mosaic of the related images for that experiment. Each measurement was taken following the subtraction of background fluorescence.

### Rationale for colocalization method used

Automated colocalization methods are suspect when applied in most systems since they require setting artificial thresholds. In our organism, the presence of the intestinal granules creates large regions of autofluorescence. We acquired the IMARIS system, which creates surface renderings of the puncta, and allowed us to remove autofluorescence, using different thresholding and machine learning protocols. We compared intensity based and pixel-classification-based surface renderings and chose intensity-based rendering.

To calculate colocalization of red and green channels, we used the algorithm object-to-object to generate 3D surfaces rendered based on signal intensity in the red, green and blue channels.

We then used machine learning to train the software to remove red and green surfaces that overlapped with the blue channel. The red and green surfaces were then analyzed for colocalization based on object-to-object statistics, specifically looking at the overlap volume ratio to surface and shortest distance to surface (surface edge to surface edge overlap). We report the % overlap for each channel out of all visible objects in that channel (Figures 2, 5). Due to the autofluorescence subtraction, the colocalization measured here may undercount how much mNG::WSHC-5 or Cadherin/HMR-1 is enriched at endosomes.

**Figure 5.**
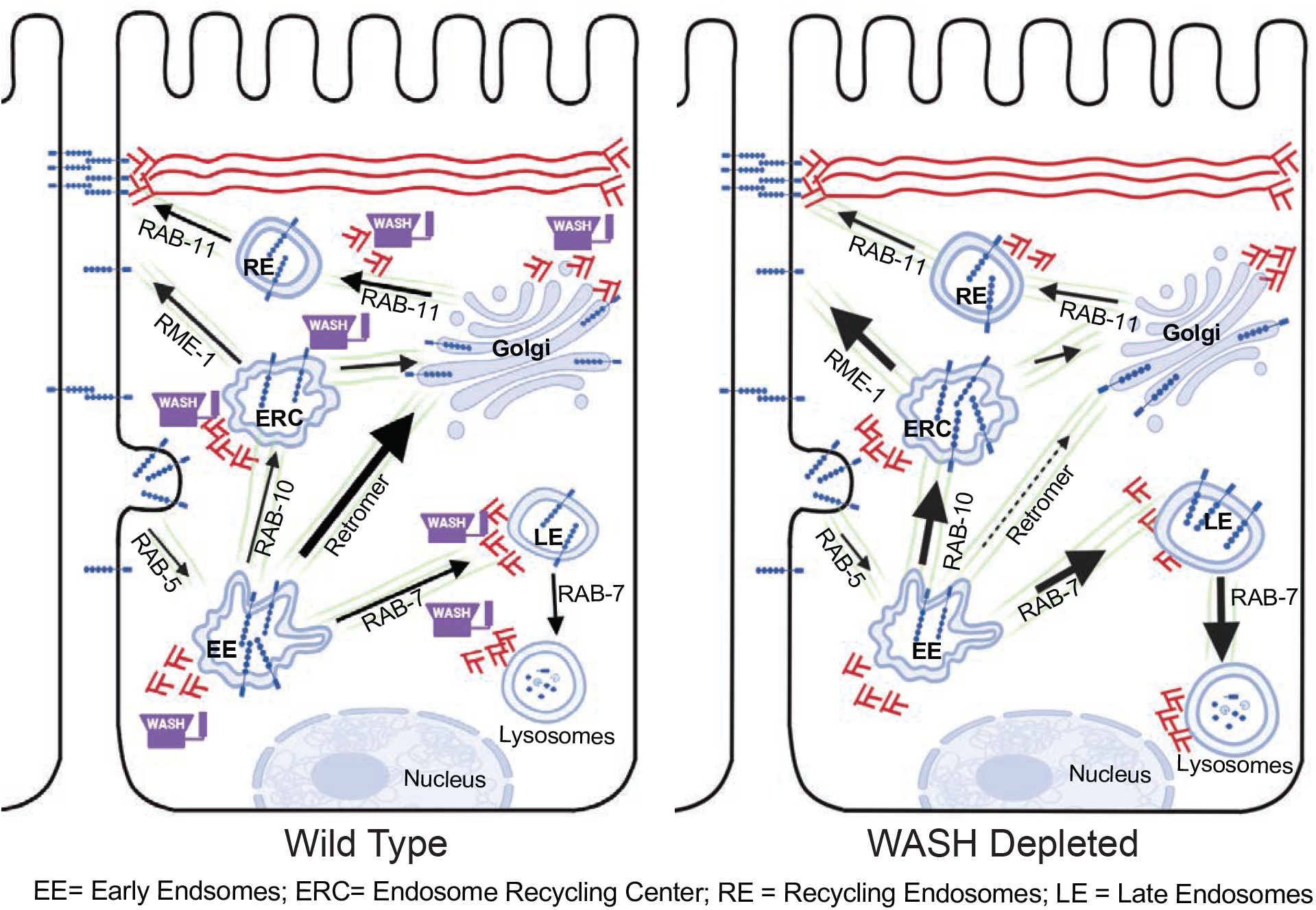
WASH regulates HMR-1/E-Cadherin transport. Model summarizing the observed changes in the sorting of Cadherin cargo in controls and in animals depleted of WASH components. Loss of WASH components led to enlarged early endosomes, and increased transport of Cadherin/HMR-1::GFP to RAB-7-positive late endosomes, and to RME-1/EHD-1-positive recycling endosomes.

### Statistical Analysis

For grouped data, statistical significance was established by performing a one-way Analysis of Variance (ANOVA), the Brown-Forysythe and Welch ANOVA, followed by a Dunnett’s multiple comparisons T3 post-test. For ungrouped data in other Figures, an unpaired t-test, the unequal variance (Welch) t test, was used. Error bars show 95% confidence intervals. Asterisks (*) denote p values *= p<.05, ** = p<0.001, *** = p<0.0001, ****=p<0.00001. All statistical analysis was performed using GraphPad Prism 8.

## Acknowledgments

We thank the NCRR-funded *Caenorhabditis* Genetics center (CGC), funded by NIH Office of Research Infrastructure Programs (P40 OD010440), Bob Goldstein and Pu Zhang for the mNG::WSHC-5 strain before publication. We thank members of the Soto Lab, Barth Grant and Mikael Garabedian for comments. This research was funded by a grant from the National Institutes of Health (NIH) (GM081670) to M.C.S., and used a Spinning Disk Microscope acquired through an NIH Shared Instrumentation Grant (1S10OD010572) to M.C.S.

---

**Fig. S1.**
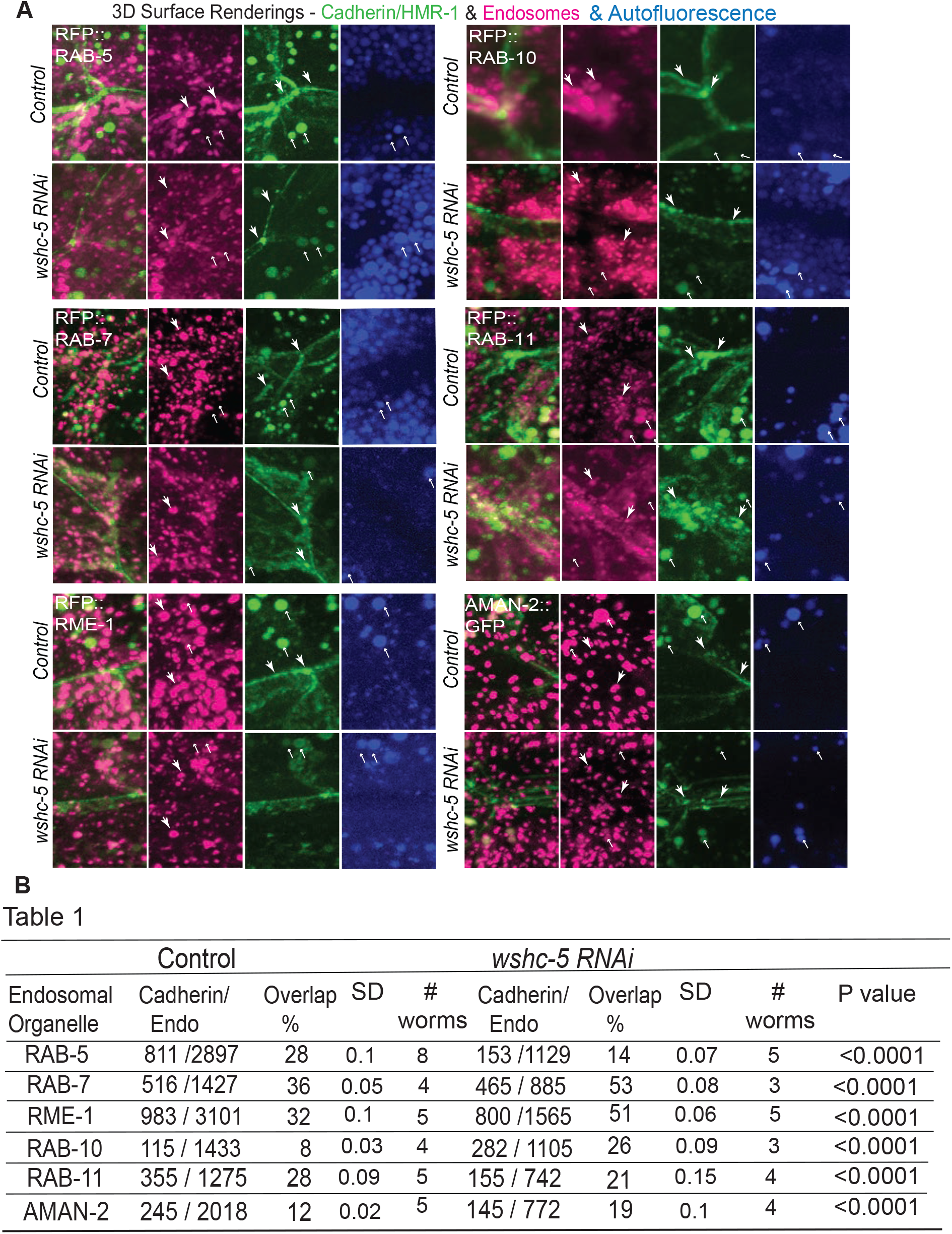
IMARIS surface renderings from raw data and raw numbers used for graph, Fig. 3D. (A) The *C. elegans* intestine is rich in autofluorescent lysosomal structures which can affect all channels, most visible in the DAPI/blue channel, labeled here Autofluorescence. Therefore, the DAPI signal was rendered in IMARIS and subtracted from the other channels, magenta and green, before applying colocalization analysis. The larger downward arrows point to sample regions where the signal is not seen in the DAPI channel, and was included in the analysis. Small upward pointing arrows show where DAPI signal was also visible in other channels, and was removed from the analysis. See Methods for further IMARIS 3D-rendering details. (B) The total individual instances of overlap between HMR-1::GFP or HMR-1::mKate2 and the endosome markers for controls and animals depleted of *wshc-5*, used to generate the statistics in Fig. 3D. Overlap % of the puncta in red and green channels was calculated using IMARIS (see Methods). Total numbers of puncta were used to calculate the SD (Standard Deviation) between individual animals, listed here are # worms. A paired T test, with Welch’s test, was used to generate P values.

## REFERENCES

Bai, Z., & Grant, B. D. (2015). A TOCA/CDC-42/PAR/WAVE functional module required for retrograde endocytic recycling. Proc Natl Acad Sci U S A, 112(12), E1443–1452. 10.1073/pnas.1418651112

Benesch, S., Polo, S., Lai, F. P., Anderson, K. I., Stradal, T. E., Wehland, J., & Rottner, K. (2005). N-WASP deficiency impairs EGF internalization and actin assembly at clathrin-coated pits. J Cell Sci, 118(Pt 14), 3103–3115. 10.1242/jcs.02444

Brüser, L., & Bogdan, S. (2017). Adherens Junctions on the Move-Membrane Trafficking of E-Cadherin. Cold Spring Harb Perspect Biol, 9(3). 10.1101/cshperspect.a029140

Campellone, K. G., Webb, N. J., Znameroski, E. A., & Welch, M. D. (2008). WHAMM is an Arp2/3 complex activator that binds microtubules and functions in ER to Golgi transport. Cell, 134(1). 10.1016/j.cell.2008.05.032

Cordova-Burgos, L., Patel, F. B., & Soto, M. C. (2021). E-Cadherin/HMR-1 Membrane Enrichment Is Polarized by WAVE-Dependent Branched Actin. Journal of Developmental Biology, 9(2), 19. 10.3390/jdb9020019

Cordova-Burgos, L., Rao, D., Egwuonwu, J., Borinskaya, S., Sasidharan, S., & Soto, M. (2023). WAVE facilitates polarized E-cadherin transport. Mol Biol Cell, 34(5), ar44. 10.1091/mbc.E22-08-0322

Chen, C. C., Schweinsberg, P. J., Vashist, S., Mareiniss, D. P., Lambie, E. J., & Grant, B. D. (2006). RAB-10 is required for endocytic recycling in the Caenorhabditis elegans intestine. Mol Biol Cell, 17(3), 1286–1297. 10.1091/mbc.e05-08-0787

Chiang, W. C., Ching, T. T., Lee, H. C., Mousigian, C., & Hsu, A. I. (2012). HSF-1 regulators DDL-1/2 link insulin-like signaling to heat-shock responses and modulation of longevity. Cell, 148(1-2). 10.1016/j.cell.2011.12.019

Derivery, E., Sousa, C., Gautier, J. J., Lombard, B., Loew, D., & Gautreau. (2009). The Arp2/3 activator WASH controls the fission of endosomes through a large multiprotein complex. Developmental cell, 17(5). 10.1016/j.devcel.2009.09.010

Duleh, S.N., & Welch, M. D. (2010). WASH and the Arp2/3 complex regulate endosome shape and trafficking. Cytoskeleton (Hoboken, N.J.), 67(3). 10.1002/cm.20437

Engqvist-Goldstein, A. E., & Drubin, D. G. (2003). Actin assembly and endocytosis: from yeast to mammals. Annu Rev Cell Dev Biol, 19, 287–332. 10.1146/annurev.cellbio.19.111401.093127

Fokin, A. I., David, V., Oguievetskaia, K., Derivery, E., Stone, C. E., Cao, L., Rocques, N., Molinie, N., Henriot, V., Aumont-Nicaise, M., Hinckelmann, M. V., Saudou, F., Le Clainche, C., Carter, A. P., Romet-Lemonne, G., & Gautreau, A. M. (2021). The Arp1/11 minifilament of dynactin primes the endosomal Arp2/3 complex. Sci Adv, 7(3). 10.1126/sciadv.abd5956

Giulianai, C., Troglio, F., Bai, Z., Patel, F. B., Zucconi, A., Malabarba, M. G., Disanza, A., Stradal, T. B., Cassata, G., Confalonieri, S., Hardin, J. D., Soto, M. C., Grant, B. D., & Scita. (2009). Requirements for F-BAR proteins TOCA-1 and TOCA-2 in actin dynamics and membrane trafficking during Caenorhabditis elegans oocyte growth and embryonic epidermal morphogenesis. PLOS Genetics, 5(10). 10.1371/journal.pgen.1000675

Gleason, A. M., Nguyen, K. C., Hall, D. H., & Grant, B. D. (2016). Syndapin/SDPN-1 is required for endocytic recycling and endosomal actin association in the C. elegans intestine. Mol Biol Cell, 27(23), 3746–3756. 10.1091/mbc.E16-02-0116

Gomez, T. S., & Billadeau, D. D. (2009). A FAM21-containing WASH complex regulates retromer-dependent sorting. Developmental cell, 17(5). 10.1016/j.devcel.2009.09.009

Hansen, M., Hsu, A. L., Dillin, A., & Kenyon, C. (2005). New genes tied to endocrine, metabolic, and dietary regulation of lifespan from a Caenorhabditis elegans genomic RNAi screen. PLoS Genet, 1(1), 119–128. 10.1371/journal.pgen.0010017

Hernandez-Valladares, M., Kim, T., Kannan, B., Tung, A., Aguda, A. H., Larsson, M., Cooper, J. A., & Robinson, R. C. (2010). Structural characterization of a capping protein interaction motif defines a family of actin filament regulators. Nat Struct Mol Biol, 17(4), 497–503. 10.1038/nsmb.1792

Hierro, A., Rojas, A. L., Rojas, R., Murthy, N., Effantin, G., Kajava, A. V., Steven, A. C., Bonifacino, J. S., & Hurley, J. H. (2007). Functional architecture of the retromer cargo-recognition complex. Nature, 449(7165), 1063–1067. 10.1038/nature06216

Jia, D., Gomez, T. S., Billadeau, D. D., & Rosen, M. K. (2012). Multiple repeat elements within the FAM21 tail link the WASH actin regulatory complex to the retromer. Mol Biol Cell, 23(12), 2352– 2361. 10.1091/mbc.E11-12-1059

Jia, D., Gomez, T. S., Metlagel, Z., Umetani, J., Otwinowski, Z., Rosen, M. K., & Billadeau, D. D. (2010). WASH and WAVE actin regulators of the Wiskott-Aldrich syndrome protein (WASP) family are controlled by analogous structurally related complexes. Proc Natl Acad Sci U S A, 107(23), 10442–10447. 10.1073/pnas.0913293107

Liu, R., Abreu-Blanco, M. T., Barry, K. C., Linardopoulou, E. V., Osborn, G. E., & Parkhurst, S. M. (2009). Wash functions downstream of Rho and links linear and branched actin nucleation factors. Development (Cambridge, England), 136(16). 10.1242/dev.035246

Marston, D. J., Higgins, C. D., Peters, K. A., Cupp, T. D., Dickinson, D. J., Pani, A. M., Moore, R. P., Cox, A. H., Kiehart, D. P., & Goldstein, B. (2016). MRCK-1 Drives Apical Constriction in C. elegans by Linking Developmental Patterning to Force Generation. Current biology : CB, 26(16), 2079–2089. 10.1016/j.cub.2016.06.010

Matthew, N. J. S., Division of, C., Molecular, M., Howard Hughes Medical Institute, U. o. C. a. S. D. S. o. M. L. J. C., McCaffery, J. M., Division of, C., Molecular, M., Howard Hughes Medical Institute, U. o. C. a. S. D. S. o. M. L. J. C., Scott, D. E., Division of, C., Molecular, M., & Howard Hughes Medical Institute, U. o. C. a. S. D. S. o. M. L. J. C. (1998). A Membrane Coat Complex Essential for Endosome-to-Golgi Retrograde Transport in Yeast. The Journal of Cell Biology, 142(3), 665. 10.1083/jcb.142.3.665

Mooren, O. L., Stuchell-Brereton, M. D., McConnell, P., Yan, C., Wilkerson, E. M., Goldfarb, D., Cooper, J. A., Sept, D., & Soranno, A. (2023). Biophysical Mechanism of Allosteric Regulation of Actin Capping Protein. J Mol Biol, 435(24), 168342. 10.1016/j.jmb.2023.168342

Morris, C., Foster, O. K., Handa, S., Peloza, K., Voss, L., Somhegyi, H., Jian, Y., Vo, M. V., Harp, M., Rambo, F. M., Yang, C., & Hermann, G. J. (2018). Function and regulation of the Caenorhabditis elegans Rab32 family member GLO-1 in lysosome-related organelle biogenesis. PLOS Genetics, 14(11). 10.1371/journal.pgen.1007772

Patel, F. B., & Soto, M. C. (2013). WAVE/SCAR promotes endocytosis and early endosome morphology in polarized C. elegans epithelia. Dev Biol, 377(2), 319–332. 10.1016/j.ydbio.2013.03.012

Romano-Moreno, M., Astorga-Simón, E. N., Rojas, A. L., & Hierro. (2024). Retromer-mediated recruitment of the WASH complex involves discrete interactions between VPS35, VPS29, and FAM21. Protein science : a publication of the Protein Society, 33(5). 10.1002/pro.4980

Shi, A., Liu, O., Koenig, S., Banerjee, R., Chen, C.C., Eimer, S., and Grant, B.D. 2012. RAB-10-GTPase-mediated regulation of endosomal phosphatidylinositol-4,5-bisphosphate. PNAS 2306–2315. 10.1073/pnas.1205278109

Smolyn, Jennifer (2020) “The first characterization of the WASH complex in C. elegans endocytic recycling,” Master’s Thesis, Rutgers University. https://rucore.libraries.rutgers.edu/rutgers-lib/62566/record/

Thomas, B. J., Wight, I. E., Chou, W. Y. Y., Moreno, M., Dawson, Z., Homayouni, A., Huang, H., Kim, H., Jia, H., Buland, J. R., Wambach, J. A., Cole, F. S., Pak, S. C., Silverman, G. A., & Luke, C. J. (2019). CemOrange2 fusions facilitate multifluorophore subcellular imaging in C. elegans. PLoS One, 14(3), e0214257. 10.1371/journal.pone.0214257

Treusch, S., Knuth, S., Slaugenhaupt, S. A., Goldin, E., Grant, B. D., & Fares, H. (2004). Caenorhabditis elegans functional orthologue of human protein h-mucolipin-1 is required for lysosome biogenesis. Proc Natl Acad Sci U S A, 101(13), 4483–4488. 10.1073/pnas.0400709101

Velle, K.B., Swafford, A.J.M., Garner, E., Fritz-Laylin, L.K. (2024). Actin network evolution as a key driver of eukaryotic diversification. J Cell Sci, 137 (15): jcs261660. 10.1242/jcs.261660

Woichansky, I., Beretta, C. A., Berns, N., & Riechmann, V. (2016). Three mechanisms control E-cadherin localization to the zonula adherens. Nat Commun, 7, 10834. 10.1038/ncomms10834

Zhang, P., Medwig-Kinney, T. N., Breiner, E. A., Perez, J. M., Song, A. N., & Goldstein. (2025). Cell signaling facilitates apical constriction by basolaterally recruiting Arp2/3 via Rac and WAVE. The Journal of Cell Biology, 224(5). 10.1083/jcb.202409133

